# CBP60-DB: An AlphaFold-predicted plant kingdom-wide database of the CALMODULIN-BINDING PROTEIN 60 (CBP60) protein family with a novel structural clustering algorithm

**DOI:** 10.1101/2022.07.07.499200

**Authors:** Keaun Amani, Vanessa Shivnauth, Christian Danve M. Castroverde

## Abstract

Molecular genetic analyses in the model species *Arabidopsis thaliana* have demonstrated the major roles of different CAM-BINDING PROTEIN 60 (CBP60) proteins in growth, stress signaling, and immune responses. Prominently, CBP60g and SARD1 are paralogous CBP60 transcription factors that regulate numerous components of the immune system, such as cell surface and intracellular immune receptors, MAP kinases, WRKY transcription factors, and biosynthetic enzymes for immunity-activating metabolites salicylic acid (SA) and *N*-hydroxypipecolic acid (NHP). However, their function, regulation and diversification in most species remain unclear. Here we have created CBP60-DB, a structural and bioinformatic database that comprehensively characterized 1052 *CBP60* gene homologs (encoding 2376 unique transcripts and 1996 unique proteins) across 62 phylogenetically diverse genomes in the plant kingdom. We have employed deep learning-predicted structural analyses using AlphaFold2 and then generated dedicated web pages for all plant CBP60 proteins. Importantly, we have generated a novel clustering visualization algorithm to interrogate kingdom-wide structural similarities for more efficient inference of conserved functions across various plant taxa. Because well-characterized CBP60 proteins in *Arabidopsis* are known to be transcription factors with putative calmodulin-binding domains, we have integrated external bioinformatic resources to analyze protein domains and motifs. Collectively, we present a plant kingdom-wide identification of this important protein family in a user-friendly AlphaFold-anchored database, representing a novel and significant resource for the broader plant biology community.

## Introduction

Plants employ constitutive and inducible defense mechanisms to combat invading pests and pathogens (Wittstock and Gershenzon, 2002; Freeman, 2008; Zhou and Zhang, 2021). A central inducible defense response is the production of the plant hormone salicylic acid (SA), which has essential roles in immunity (Ding & Ding, 2020; Peng et al., 2021; Shields et al., 2022) and abiotic stress tolerance (Gharbi et al., 2018; Khan et al., 2019; Saleem et al., 2021). Thorough understanding of plant immunity and stress responses are important in reducing global crop losses and ensuring food security worldwide (Bailey-Serres et al., 2019; Savary et al., 2019).

In the model plant species *Arabidopsis thaliana*, SA production in response to stress is mediated by the sequential action of the ISOCHORISMATE SYNTHASE 1 (ICS1), ENHANCED DISEASE SUSCEPTIBILITY 5 (EDS5) and AVRPPHB SUSCEPTIBLE 3 (PBS3) proteins (Rekhter et al., 2019), which are controlled at the transcriptional level by the master transcription factor CAM-BINDING PROTEIN 60-LIKE G (CBP60g) and its functionally redundant homolog SAR Deficient 1 (SARD1; Wang et al., 2009; Zhang et al., 2010; Wang et al., 2011; Sun et al., 2015). Notably, it is known that SA production and plant immunity are vulnerable to warming temperatures (Huot et al., 2017; Castroverde and Dina, 2021). This critical temperature-vulnerability of the plant immune system is controlled via CBP60g/SARD1 (Kim et al., 2022), which are members of the broadly conserved plant CBP60 protein family (Zheng et al., 2021).

Biological understanding of protein function relies on detailed characterization of protein structures. However, accurate prediction of protein structure from amino acid sequence alone has remained a central problem in biology (Dill et al., 2008). Traditional methods, such as X-ray crystallography or NMR spectroscopy, are usually very expensive, time consuming, and can fail to produce viable results for complexes, membrane-bound proteins, or proteins that are unable to crystallize (Tugarinov et al., 2004; Shi, 2014; Nogales and Scheres, 2015). A major advance to solve this grand challenge occurred with the launch of AlphaFold2, which is a novel deep learning approach for accurately predicting the three-dimensional structure of a protein from its amino acid sequence (Jumper et al., 2021). However, base AlphaFold2 also suffers from a few drawbacks, such as lack of exposure for certain internal settings (e.g., number of recycling steps), it is slightly unoptimized, and the default MSA generation algorithms used can be slow and time-consuming (Mirdita et al., 2022). ColabFold (Mirdita et al., 2022) is an AlphaFold2 derivative that addresses the aforementioned issues with AlphaFold2 and is able to generate highly accurate predictions comparable, if not superior to those of AlphaFold2. Furthermore, ColabFold can produce more predictions within a shorter period.

Because of the biological importance of CBP60g and SARD1 proteins for plant immune system resilience under changing environmental conditions (Wan et al., 2012; Choudhary et al., 2022; Kim et al., 2022), it is critical that we fully understand their structures and functions in other plants. This mechanistic knowledge has critical ramifications on safeguarding plant disease resistance for a warming climate. Although a recent study conducted a kingdom-wide phylogenetic analysis of the CBP60 family and potential protein neofunctionalization (Zheng et al., 2021), there is little functional and molecular information on these proteins in most plants, including agriculturally important crop species.

To further understand the diversity of CBP60 protein structure and function in the plant kingdom, we have created a fully curated, AlphaFold-generated (Jumper et al., 2021) structural database called the Plant CBP60 Protein Family Database or CBP60-DB (https://cbp60db.wlu.ca/). Of note, this paper describes an algorithm that to our knowledge is a novel approach to accurately clustering proteins by structural similarity. The proposed algorithm is simple, accurate, and can be easily reproduced on any modern device. By building our novel visual clustering algorithm, we were able to compare and cluster the predicted protein structures, facilitating easier ortholog selection and inference of putative biological functions. A Google Colaboratory notebook is provided, as well as a minimal implementation for executing locally. We have showcased a visualization for this algorithm on the index page of the CBP60-DB web application.

### Plant kingdom-wide sequence collection and AlphaFold-based protein folding

We first identified CBP60 genes and proteins in plant species with published and fully sequenced genomes. Using the Gramene comparative genomics website (http://gramene.org/; Tello-Ruiz et al., 2020), we obtained a comprehensive kingdom-wide list of representative plant species and *CBP60* gene homologs in these species. Our base dataset consisted of species names, gene sequences, transcript/cDNA sequences, and protein sequence data. Each protein entry’s amino acid sequence was used as an input to ColabFold for structural predictions. ColabFold (https://github.com/sokrypton/ColabFold) was used instead of the original AlphaFold2 since the former produces a higher number of predictions within a shorter time, while also improving prediction quality compared to base AlphaFold2. This improvement is primarily due to ColabFold’s usage of the MMseqs2 algorithm for faster homology search as well as other model optimizations (Steinegger and Söding, 2017). Furthermore, ColabFold makes some of AlphaFold2’s internal settings easily accessible and configurable, allowing us to adjust settings such as the number of recycling iterations.

### Database implementation

Prior to the development of CBP60-DB and its web application components, we determined that an effective solution must be scalable, responsive, simple to use, and sufficiently modular, so that the application could easily be adapted to other protein families and similar projects. The final version of our database contains 1996 unique predicted structures (from 2376 corresponding cDNA/transcripts and 1052 unique genes), as well as corresponding metadata, and confidence metrics. The predicted structures are available in the protein databank (PDB; Berman et al., 2000) and the newer macromolecular Crystallographic Information File (mmCIF) file formats.

The CBP60-DB user interface was designed to be easy to navigate, with an emphasis on several intuitive visualization options that are available and assembled for best user accessibility. Additionally, the application was written in the Go programming language without third party dependencies, making it straightforward to re-deploy across any modern system. All database contents are either stored within the assets directory of the application, which is freely accessible via HTTP(S), or stored within an internal json file that is then loaded into memory as a hash table, where keys are the md5 hashes of the unique transcript names. The advantage of an internal hash table over a traditional database management system (DBMS) is that the internal hash table is faster for accessing and serving data and requires no additional dependencies. Furthermore, since the contents of the database are static and the memory required to load the json file is reasonable (6.9 Mb), there is little need for using an alternative DBMS. However, should we decide to scale the contents of the database to include vastly more entries, an alternative DBMS will be the preferable solution.

### Data archival

CBP60-DB archives and provides access to the following data below. Note that protein structures which have been updated, replaced, or removed will not be archived.

- Predicted protein crystal structure in PDB and mmCIF file formats.
- Protein metadata and AlphaFold2 metadata in json format.
- Generated thumbnails of the predicted structure in png format.
- AlphaFold2 scoring metrics in json format (pLDDT, PAE, and pTM score).
- MMseqs2 MSA file used during model inference in a3m format.
- Cluster map of predicted structures in json format.
- Phylogenetic tree created within MEGA using the MUltiple Sequence Comparison by Log-Expectation (MUSCLE) alignment algorithm in FASTA format.
- Phylogenetic tree generated by FastTree in the Newick file format.

The predicted Local Distance Difference Test (pLDDT-Cα) is a per residue metric used by AlphaFold2 to gauge the model’s confidence in the position and orientation of each residue within a predicted structure. Values range from 0 to 100, where higher values are associated with greater prediction accuracy and less disorder (Jumper et al., 2021).

The Predicted Aligned Error (PAE) is a **N_res_** x **N_res_** matrix where **N_res_** corresponds to the number of residues within the input amino acid sequence. Each element within the matrix represents the predicted distance error in Ångströms of the 1^st^ residue’s position when aligned on the 2^nd^ residue (Varadi et al., 2021).

The Molecular Evolutionary Genetics Analysis (Tamura et al., 2021) application was used to produce the alignment fasta file using the MUltiple Sequence Comparison by Log-Expectation (MUSCLE) (Edgar, 2004) algorithm with the following parameters (Supplementary Table 1). The alignment fasta file was then used by the Fast Tree algorithm (Price et al., 2009) to produce a phylogenetic tree in the Newick file format.

### Clustering proteins by structural similarity

Clustering proteins by their structural similarity is an invaluable method for finding proteins with potentially similar function but with diverging sequences, especially for large protein families (Mai et al., 2016; Teletin et al., 2019). Traditional sequence-based cluster algorithms also provide simple and computationally efficient ways of representing similar proteins but have a major drawback with regards to proteins with similar functionality but different sequences (Krissinel, 2007; Kosloff and Kolodny, 2007).

We have proposed a novel algorithm that is simple and effective at clustering proteins by structural similarity, while also being easily parallelizable. Our algorithm utilizes metrics used for protein structure comparison (e.g., TM-Align, Root Mean Square Deviation (RMSD), etc.) to produce a feature tensor that is then used as input to the Uniform Manifold Approximation and Projection (UMAP; McInnes et al., 2018) algorithm. The corresponding UMAP projection can then be used as an intuitive visualization, where proteins that are more structurally similar to one another will be clustered within closer proximity to each other. The advantages of our algorithm are that it is trivial to implement, easy to utilize, and highly configurable with regards to feature selection and UMAP hyper-parameter tuning. Furthermore, our algorithm can cluster small datasets of protein with minimal hardware and within a reasonable amount of time. However, a drawback of the algorithm is its quadratic time complexity which does not allow it to efficiently scale on lower-end hardware.

To produce the input feature tensor pairwise structural comparison, metrics such as TM-Align (Zhang, 2005) optionally alongside other metrics such as RMSD were used to produce a □ × □ × □ feature tensor, where □ is the number of proteins and □ is the number of features. The feature tensor was then flattened to produce a □ x (□ x □) matrix, which was used as an input to UMAP. UMAP is a powerful dimensionality reduction algorithm that can generally create more meaningful representations compared to principal component analysis, while also outperforming t-distributed stochastic neighbor embedding (t-SNE; McInnes et al., 2018). It is also noteworthy to mention that swapping UMAP with t-SNE produces comparable projections; however, UMAP is significantly faster and, in our opinion, generally produces more intuitive projections.

A Google Colaboratory notebook demo (https://colab.research.google.com/drive/1LOZY33CSO5-PdJAdDApyPlfxUu4DHjcW) and minimal Python implementation for the clustering algorithm are available. Additionally, a structural cluster of all proteins available within the CBP60-DB is available on the index page of the application (Fig 1).

**Fig 1.**
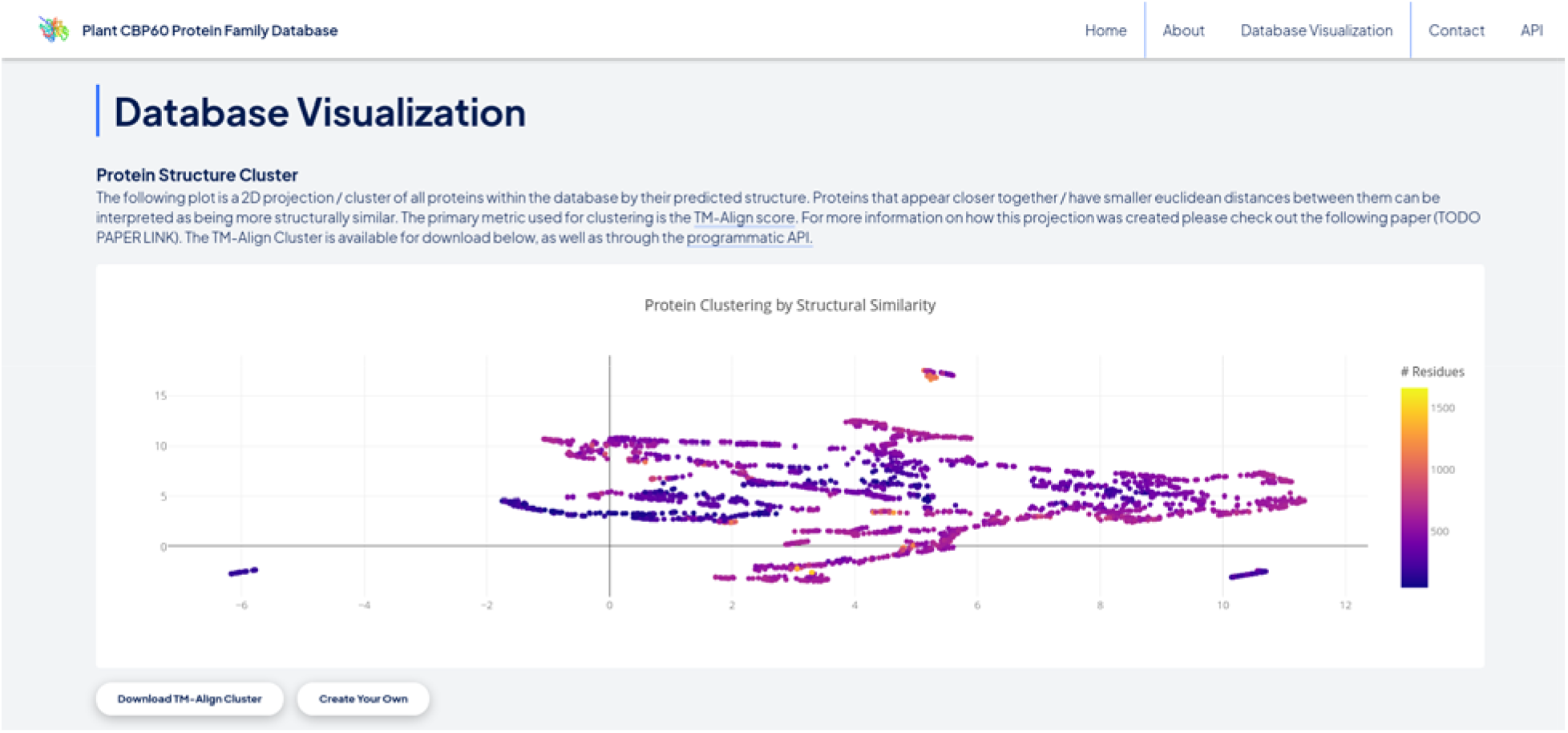
Screenshot of the top of the protein structure cluster of the entire CBP60-DB within the Database Visualization section of the index page.

### Data access

Information stored on CBP60-DB is hosted by Wilfrid Laurier University, which can be accessed using a modern web browser (https://cbp60db.wlu.ca/), through the application programming interface (API), as well as through our public github repository (https://github.com/KeaunAmani/cbp60db/). Accessing the database through either means provides access to all data including the predicted structures, metadata, thumbnails, prediction metrics, as well as the cluster map. Additionally, viewing CBP60-DB via your web browser provides access to several interactive and intuitive visualizations, featuring an interactive protein viewer, navigable cluster map (Fig 1), interactive plots for prediction metrics, as well as the top five most structurally similar proteins (if available).

### Navigating the database

There are three primary web pages available on CBP60-DB: (1) index page, (2) protein search page, and (3) protein information page.

#### Index page

The CBP60-DB index page (Fig 2) acts as the website home page containing general information, navigation options, database visualizations, downloads, as well as API endpoint documentation. To navigate the database, users may either search for a protein directly via the search bar in the page header, or alternatively interact with the protein structural visualization cluster. By clicking a node within the cluster, users will be redirected to that protein’s information page. Alternatively, the aforementioned search bar allows users to search for protein by their transcript name, gene name, or source organism. Proteins that match the search query will be displayed in the following search page. Another visualization available within this page is an interactive phylogenetic tree explorer. Note that downloads for the TM-Align Cluster, FASTA Alignment, and Phylogenetic tree are available underneath their respective visualizations.

**Fig 2.**
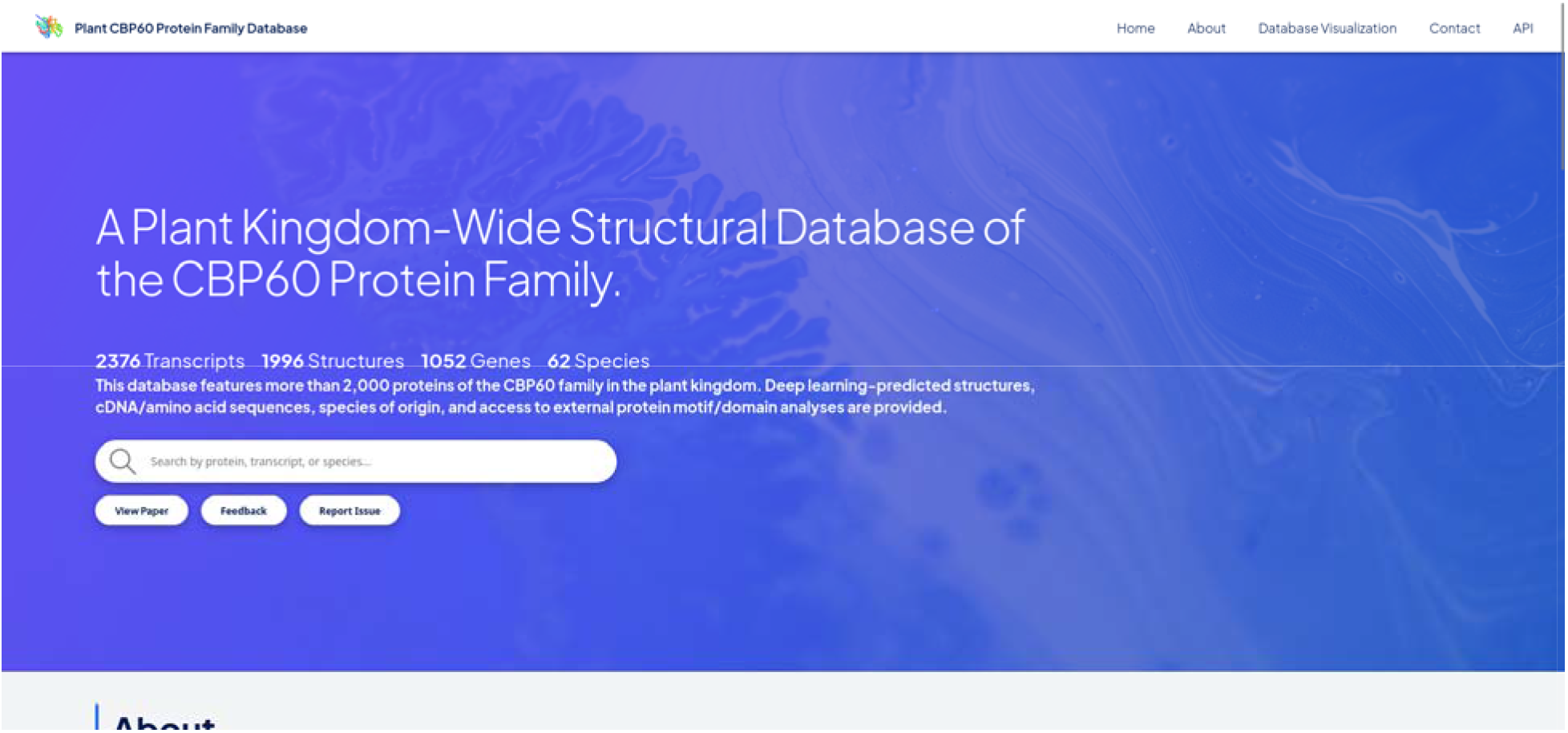
Screenshot of the top of the CBP60-DB index page.

#### Search page

The search page (Fig 3) displays the search results from queries made via the search bar on any page within CBP60-DB. Once a query is submitted through the search bar, users will be redirected to this page. If no query is provided, all database entries will be displayed instead. Search results are unordered and in the form of card previews containing a thumbnail of the predicted crystal structure, the transcript name, gene name, source organism, cDNA length, and amino acid sequence length. Users can click on cards to visit their corresponding protein information pages.

**Fig 3.**
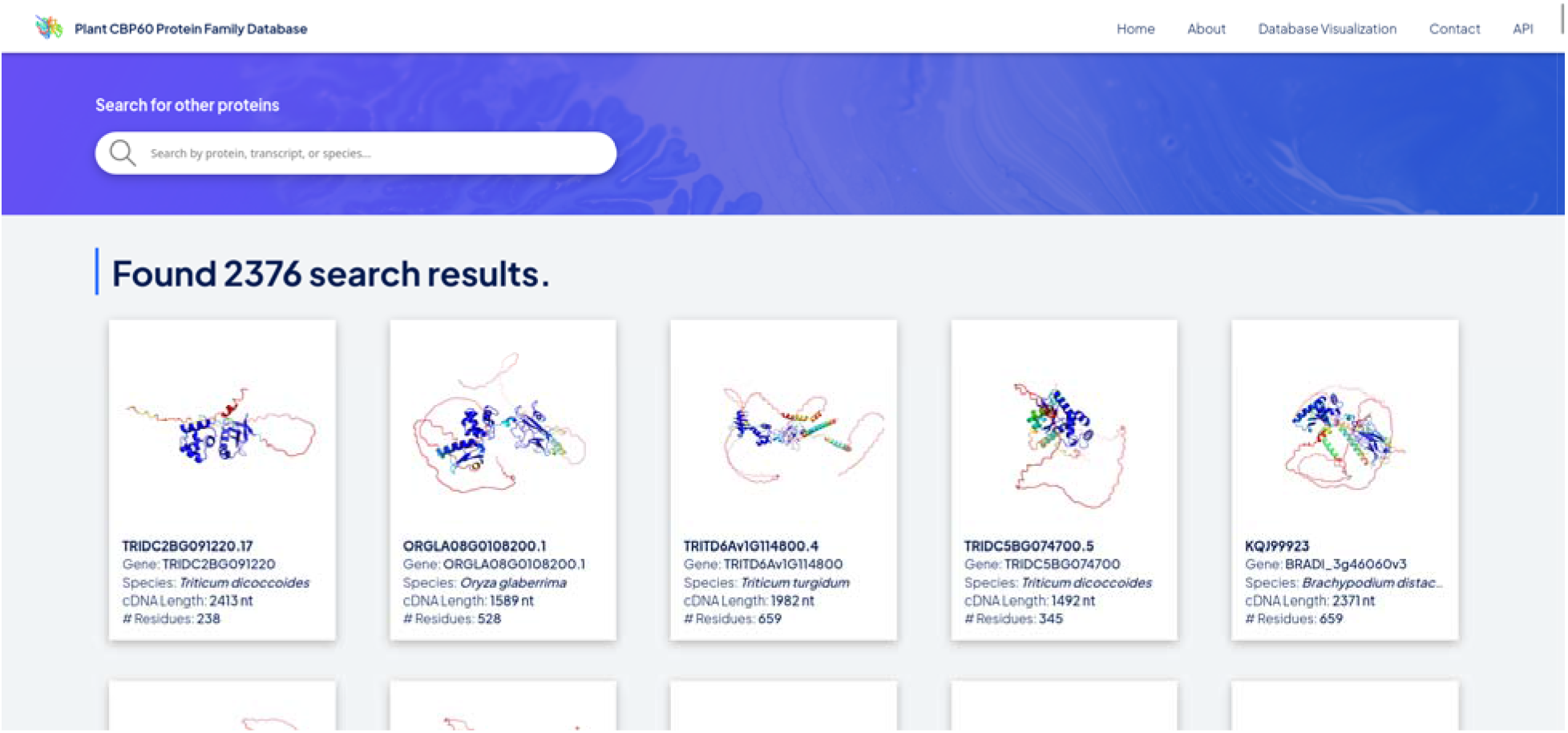
Screenshot of the top of the CBP60-DB search page.

#### Protein information page

The protein information page (Fig 4) is arguably the most useful page within CBP60-DB, providing a simple interface for viewing the gene name, transcript name, source organism, AlphaFold2 settings used, cDNA sequence, amino acid sequence, structure data, redirect to DNA-Binding Residues tool (Hwang et al., 2007), redirect to Eukaryotic Linear Motifs tool (Kumar et al., 2020), various downloads as well as visualizations, and the top five most similar structures (if available) according to the clustering algorithm.

**Fig 4.**
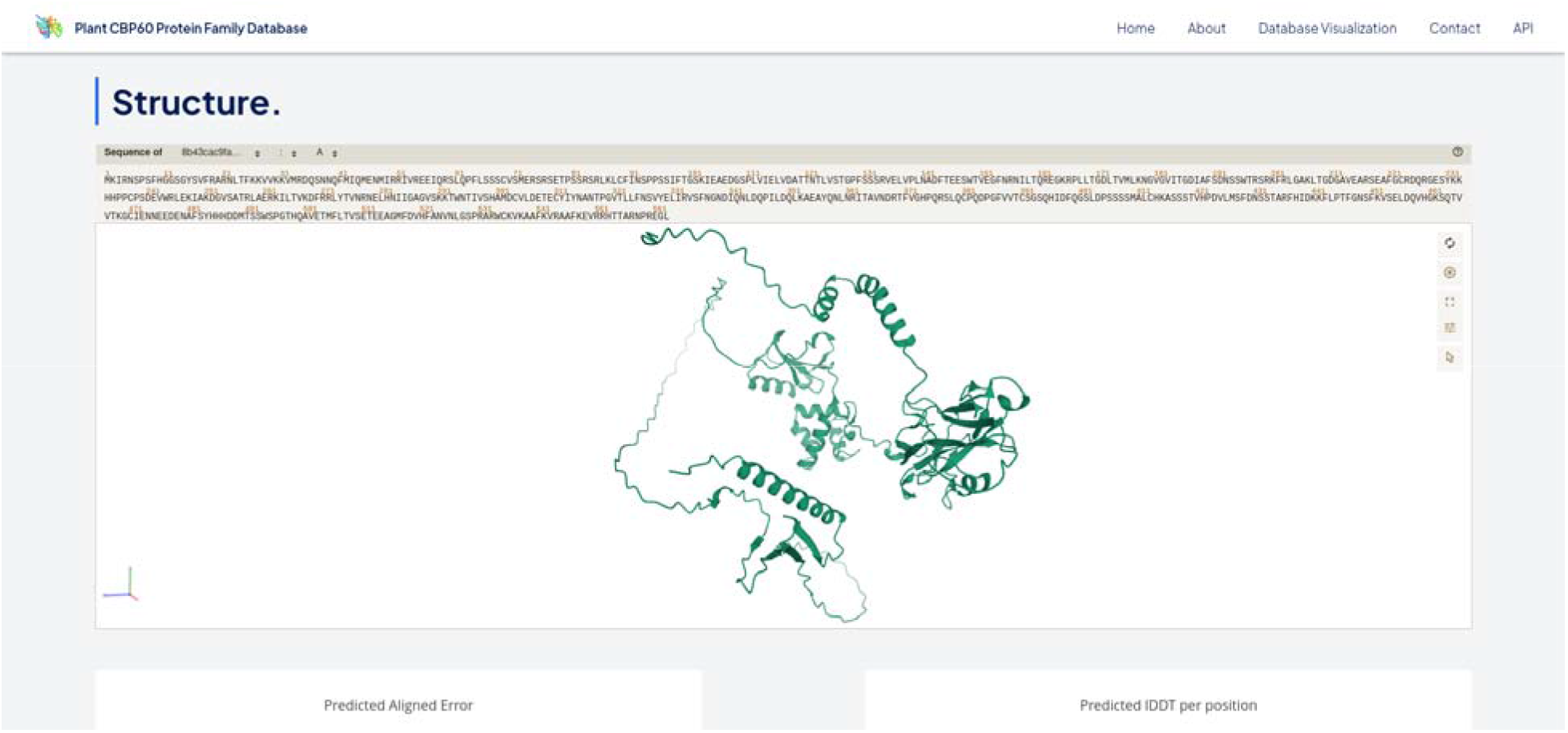
Screenshot of the top of the CBP60-DB protein information page for the representative protein with the transcript name AT5G26920.1.

Data visualizations available on this page include an interactive molecular viewer of the predicted protein structure utilizing PDBe Molstar (Sehnal et al., 2021), as well as interactive plots for the PAE and pLDDT scores powered by the plotly.js library (Plotly Technologies Inc., 2015). Additionally, the exact same protein cluster from the Index page is also available with the current protein highlighted within the plot. Similar to the Index page, this plot is also navigable in the same way.

Data downloads available on this page consist of the PDB file of the predicted structure, mmCIF file of the predicted structure, PAE json file, pLDDT json file, amino acid sequence FASTA file, and the generated MSA used to predict protein structure. These resources are also available for download directly via the programmatic API.

## Conclusion and Outlook

Recent in-silico advances for protein structure prediction have accelerated molecular biology research at an unprecedented scale. Deep learning models have now proven themselves to be effective tools for protein folding and are made even more valuable through their ease of use and lower costs compared to traditional techniques (Jumper et al., 2021; Baek et al., 2021). By determining the structures of all proteins within the CBP60 plant kingdom family, biologists can infer putative functions, evolutionary relationships, and other meaningful information from protein structures on a broader scale.

Overall, the CBP60-DB has generated useful and comprehensive datasets that are foundational for further functional and molecular studies. Because CBP60 protein family members CBP60 and SARD1 are indispensable master regulators of plant defense responses (Wang et al., 2009; Zhang et al., 2010; Wang et al., 2011; Sun et al., 2015; Kim et al., 2022), our fundamental understanding of their structural and functional diversity has profound implications for mitigating plant diseases. This could potentially address major challenges in agricultural and natural ecosystems globally, especially on understanding plant immune system resilience (Velasquez et al., 2018; Kim et al., 2021; Kim et al., 2022) to boost worldwide crop productivity (Bailey-Serres et al., 2019). Using a robust and rapid bioinformatic pipeline, our comprehensive deep learning-assisted database with a novel structural clustering algorithm provides the scientific community with easy-to-access candidate genes/proteins that can be further engineered to strengthen plant health in a changing world.

## Acknowledgements

Research in the Castroverde Lab is funded by the Natural Sciences and Engineering Research Council of Canada (NSERC) Discovery Grant (to C.D.M.C.), Canada Foundation for Innovation (to C.D.M.C.), Ontario Research Fund (to C.D.M.C.), Laurier Faculty of Science institutional start-up funds (to C.D.M.C.), and a Mitacs Research Training Award (to V.S.). We also thank Compute Canada and SHARCNET for providing computational power and support, Laurier Information and Communication Technologies for website hosting, as well as Dr. Sean Johnson for his critical guidance and feedback.

---

**Supplementary Table 1:** Configuration used for MUSCLE alignment generation using MEGA11.

| Option | Value used |
| --- | --- |
| Gap Open Penalty | -2.90 |
| Gap Extend Penalty | 0.00 |
| Hydrophobicity Multiplier Penalty | 1.20 |
| Max Memory in MB | 4096 |
| Max Iterations | 16 |
| Cluster Method (Iterations 1,2) | UPGMA |
| Cluster Method (Other Iterations) | UPGMA |
| Min Diag Length (Lambda) | 24 |

## References

1. Ahdritz G, Bouatta N, Kadyan S, Xia Q, Gerecke W, AlQuraishi M. 2021. OpenFold. doi:10.5281/zenodo.5709539. https://github.com/aqlaboratory/openfold.

2. Baek M, DiMaio F, Anishchenko I, Dauparas J, Ovchinnikov S, Lee GR, Wang J, Cong Q, Kinch LN, Schaeffer RD, et al. 2021. Accurate prediction of protein structures and interactions using a three-track neural network. Science. 373(6557):871–876. doi:10.1126/science.abj8754.

3. Bailey-Serres J, Parker JE, Ainsworth EA, Oldroyd GED, Schroeder JI. 2019. Genetic strategies for improving crop yields. Nature. 575(7781):109–118. doi:10.1038/s41586-019-1679-0. https://www.nature.com/articles/s41586-019-1679-0.

4. Berman HM. 2000. The Protein Data Bank. Nucleic Acids Research. 28(1):235–242. doi:10.1093/nar/28.1.235.

5. Castroverde CDM, Dina D. 2021. Temperature regulation of plant hormone signaling during stress and development. Journal of Experimental Botany. doi:10.1093/jxb/erab257.

6. Cheng S, Wu R, Yu Z, Li B, Zhang X, Peng J, You Y. 2022. FastFold: Reducing AlphaFold Training Time from 11 Days to 67 Hours. arXiv:220300854 [cs, q-bio]. https://arxiv.org/abs/2203.00854.

7. Choudhary A, Senthil Kumar M. 2022. Drought attenuates plant defence against bacterial pathogens by suppressing the expression of *CBP60g* / *SARD1* during combined stress. Plant, Cell & Environment. 45(4):1127–1145. doi:10.1111/pce.14275.

8. Dill KA, Ozkan SB, Shell MS, Weikl TR. 2008. The Protein Folding Problem. Annual Review of Biophysics. 37(1):289–316. doi:10.1146/annurev.biophys.37.092707.153558.

9. Ding P, Ding Y. 2020 Feb. Stories of Salicylic Acid: A Plant Defense Hormone. Trends in Plant Science. doi:10.1016/j.tplants.2020.01.004.

10. Edgar RC. 2004. MUSCLE: multiple sequence alignment with high accuracy and high throughput. Nucleic Acids Research. 32(5):1792–1797. doi:10.1093/nar/gkh340.

11. Freeman. 2008. An Overview of Plant Defenses against Pathogens and Herbivores. The Plant Health Instructor. doi:10.1094/phi-i-2008-0226-01.

12. Gharbi E, Lutts S, Dailly H, Quinet M. 2018. Comparison between the impacts of two different modes of salicylic acid application on tomato (Solanum lycopersicum) responses to salinity. Plant Signaling & Behavior. 13(5):e1469361. doi:10.1080/15592324.2018.1469361.

13. Huot B, Castroverde CDM, Velásquez AC, Hubbard E, Pulman JA, Yao J, Childs KL, Tsuda K, Montgomery BL, He SY. 2017. Dual impact of elevated temperature on plant defence and bacterial virulence in Arabidopsis. Nature Communications. 8(1). doi:10.1038/s41467-017-01674-2.

14. Hwang S, Gou Z, Kuznetsov IB. 2007. DP-Bind: a web server for sequence-based prediction of DNA-binding residues in DNA-binding proteins. Bioinformatics. 23(5):634–636. doi:10.1093/bioinformatics/btl672.

15. Jumper J, Evans R, Pritzel A, Green T, Figurnov M, Ronneberger O, Tunyasuvunakool K, Bates R, Žídek A, Potapenko A, et al. 2021. Highly accurate protein structure prediction with AlphaFold. Nature. 596(7873):583–589. doi:10.1038/s41586-021-03819-2.

16. Khan A, Kamran M, Imran M, Al-Harrasi A, Al-Rawahi A, Al-Amri I, Lee I-J, Khan AL. 2019. Silicon and salicylic acid confer high-pH stress tolerance in tomato seedlings. Scientific Reports. 9(1). doi:10.1038/s41598-019-55651-4.

17. Kim JH, Castroverde CDM, Huang S, Li C, Hilleary R, Seroka A, Sohrabi R, Medina-Yerena D, Huot B, Wang J, et al. 2022 Jun 29. Increasing the resilience of plant immunity to a warming climate. Nature. doi:10.1038/s41586-022-04902-y.

18. Kim JH, Hilleary R, Seroka A, He SY. 2021. Crops of the future: building a climate-resilient plant immune system. Current Opinion in Plant Biology. 60:101997. doi:10.1016/j.pbi.2020.101997.

19. Kosloff M, Kolodny R. 2007. Sequence-similar, structure-dissimilar protein pairs in the PDB. Proteins: Structure, Function, and Bioinformatics. 71(2):891–902. doi:10.1002/prot.21770.

20. Krissinel E. 2007. On the relationship between sequence and structure similarities in proteomics. Bioinformatics. 23:717–723. doi:10.1093/bioinformatics/btm006.

21. Kumar M, Gouw M, Michael S, Sámano-Sánchez H, Pancsa R, Glavina J, Diakogianni A, Valverde JA, Bukirova D, Čalyševa J, et al. 2020. ELM-the eukaryotic linear motif resource in 2020. Nucleic Acids Research. 48(D1):D296–D306. doi:10.1093/nar/gkz1030.

22. Mai T-L, Hu G-M, Chen C-M. 2016. Visualizing and Clustering Protein Similarity Networks: Sequences, Structures, and Functions. Journal of Proteome Research. 15(7):2123–2131. doi:10.1021/acs.jproteome.5b01031.

23. McInnes L, Healy J, Melville J. 2018. UMAP: Uniform Manifold Approximation and Projection for Dimension Reduction. arXivorg. https://arxiv.org/abs/1802.03426.

24. Mirdita M, Schütze K, Moriwaki Y, Heo L, Ovchinnikov S, Steinegger M. 2022. ColabFold: making protein folding accessible to all. Nature Methods.:1–4. doi:10.1038/s41592-022-01488-1.

25. Nogales E, Scheres Sjors HW. 2015. Cryo-EM: A Unique Tool for the Visualization of Macromolecular Complexity. Molecular Cell. 58(4):677–689. doi:10.1016/j.molcel.2015.02.019.

26. Peng Y, Yang J, Li X, Zhang Y. 2021. Salicylic Acid: Biosynthesis and Signaling. Annual Review of Plant Biology. 72:761–791. doi:10.1146/annurev-arplant-081320-092855.

27. Plotly Technologies Inc. 2015. Collaborative data science. https://plot.ly.

28. Price MN, Dehal PS, Arkin AP. 2009. FastTree: Computing Large Minimum Evolution Trees with Profiles instead of a Distance Matrix. Molecular Biology and Evolution. 26(7):1641–1650. doi:10.1093/molbev/msp077.

29. Rekhter D, Lüdke D, Ding Y, Feussner K, Zienkiewicz K, Lipka V, Wiermer M, Zhang Y, Feussner I. 2019. Isochorismate-derived biosynthesis of the plant stress hormone salicylic acid. Science. 365(6452):498–502. doi:10.1126/science.aaw1720.

30. Saleem M, Fariduddin Q, Castroverde CDM. 2021. Salicylic acid: A key regulator of redox signalling and plant immunity. Plant Physiology and Biochemistry. 168:381–397. doi:10.1016/j.plaphy.2021.10.011.

31. Savary S, Willocquet L, Pethybridge SJ, Esker P, McRoberts N, Nelson A. 2019. The global burden of pathogens and pests on major food crops. Nature Ecology & Evolution. 3(3):430–439. doi:10.1038/s41559-018-0793-y.

32. Sehnal D, Bittrich S, Deshpande M, Svobodová R, Berka K, Bazgier V, Velankar S, Burley SK, Koča J, Rose AS. 2021. Mol* Viewer: modern web app for 3D visualization and analysis of large biomolecular structures. GitHub. https://github.com/molstar/molstar.

33. Shi Y. 2014. A Glimpse of Structural Biology through X-Ray Crystallography. Cell. 159(5):995–1014. doi:10.1016/j.cell.2014.10.051.

34. Shields A, Shivnauth V, Castroverde CDM. 2022. Salicylic Acid and N-Hydroxypipecolic Acid at the Fulcrum of the Plant Immunity-Growth Equilibrium. Frontiers in Plant Science. 13. doi:10.3389/fpls.2022.841688.

35. Steinegger M, Söding J. 2017. MMseqs2 enables sensitive protein sequence searching for the analysis of massive data sets. Nature Biotechnology. 35(11):1026–1028. doi:10.1038/nbt.3988.

36. Sun T, Zhang Y, Li Y, Zhang Q, Ding Y, Zhang Y. 2015. ChIP-seq reveals broad roles of SARD1 and CBP60g in regulating plant immunity. Nature Communications. 6(1). doi:10.1038/ncomms10159.

37. Tamura K, Stecher G, Kumar S. 2021. MEGA11: Molecular evolutionary genetics analysis version 11. Molecular Biology and Evolution. 38:3022–3027. doi:10.1093/molbev/msab120.

38. Teletin M, Czibula G, Bocicor M-I. 2019. Using clustering models for uncovering proteins’ structural similarity. IEEE Xplore.:185–190. doi:10.1109/SACI46893.2019.9111642. https://ieeexplore.ieee.org/document/9111642.

39. Tello-Ruiz MK, Naithani S, Gupta P, Olson A, Wei S, Preece J, Jiao Y, Wang B, Chougule K, Garg P, et al. 2020. Gramene 2021: harnessing the power of comparative genomics and pathways for plant research. Nucleic Acids Research. 49(D1):D1452–D1463. doi:10.1093/nar/gkaa979.

40. Tugarinov V, Hwang PM, Kay LE. 2004. Nuclear Magnetic Resonance Spectroscopy of High-Molecular-Weight Proteins. Annual Review of Biochemistry. 73(1):107–146. doi:10.1146/annurev.biochem.73.011303.074004.

41. Varadi M, Anyango S, Deshpande M, Nair S, Natassia C, Yordanova G, Yuan D, Stroe O, Wood G, Laydon A, et al. 2021. AlphaFold Protein Structure Database: massively expanding the structural coverage of protein-sequence space with high-accuracy models. Nucleic Acids Research. doi:10.1093/nar/gkab1061.

42. Wan D, Li R, Zou B, Zhang X, Cong J, Wang R, Xia Y, Li G. 2012. Calmodulin-binding protein CBP60g is a positive regulator of both disease resistance and drought tolerance in Arabidopsis. Plant Cell Reports. 31(7):1269–1281. doi:10.1007/s00299-012-1247-7.

43. Wang L, Tsuda K, Sato M, Cohen JD, Katagiri F, Glazebrook J. 2009. Arabidopsis CaM Binding Protein CBP60g Contributes to MAMP-Induced SA Accumulation and Is Involved in Disease Resistance against Pseudomonas syringae. PLoS Pathogens. 5(2):e1000301. doi:10.1371/journal.ppat.1000301.

44. Wang L, Tsuda K, Truman W, Sato M, Nguyen LV, Katagiri F, Glazebrook J. 2011. CBP60g and SARD1 play partially redundant critical roles in salicylic acid signaling. The Plant Journal. 67(6):1029–1041. doi:10.1111/j.1365-313x.2011.04655.x.

45. Wittstock U, Gershenzon J. 2002. Constitutive plant toxins and their role in defense against herbivores and pathogens. Current Opinion in Plant Biology. 5(4):300–307. doi:10.1016/s1369-5266(02)00264-9.

46. Zhang Y. 2005. TM-align: a protein structure alignment algorithm based on the TM-score. Nucleic Acids Research. 33(7):2302–2309. doi:10.1093/nar/gki524.

47. Zhang Y, Xu S, Ding P, Wang D, Cheng YT, He J, Gao M, Xu F, Li Y, Zhu Z, et al. 2010. Control of salicylic acid synthesis and systemic acquired resistance by two members of a plant-specific family of transcription factors. Proceedings of the National Academy of Sciences. 107(42):18220–18225. doi:10.1073/pnas.1005225107.

48. Zheng Q, Majsec K, Katagiri F. 2021 Oct 5. Pathogen driven coevolution across the CBP60 plant immune regulator subfamilies confers resilience on the regulator module. New Phytologist. doi:10.1111/nph.17769.

49. Zhou J-M, Zhang Y. 2020. Plant Immunity: Danger Perception and Signaling. Cell. 181(5):978–989. doi:10.1016/j.cell.2020.04.028.

